# The hippocampus is necessary for the sleep-dependent consolidation of a task that does not require the hippocampus for initial learning

**DOI:** 10.1101/451195

**Authors:** Anna C. Schapiro, Allison G. Reid, Alexandra Morgan, Dara S. Manoach, Mieke Verfaellie, Robert Stickgold

## Abstract

During sleep, the hippocampus plays an active role in consolidating memories that depend on it for initial encoding. There are hints in the literature that the hippocampus may have a broader influence, contributing to the consolidation of memories that may not initially require the area. We tested this possibility by evaluating learning and consolidation of the motor sequence task (MST) in hippocampal amnesics and demographically matched control participants. While the groups showed similar initial learning, only controls exhibited evidence of sleep-dependent consolidation. These results demonstrate that the hippocampus can be required for normal consolidation of a task without being required for its acquisition, suggesting that the area plays a broader role in coordinating sleep-dependent memory consolidation than has previously been assumed.

## Introduction

The hippocampus plays an important and active role in memory consolidation during sleep^1^. It replays recent experiences during high frequency ripple oscillations^2^ that often co-occur with spindle events in neocortex^3-7^, which are also associated with replay of recent experience^8^. This hippocampal-cortical dialogue is thought to facilitate the transfer of new memories encoded in the hippocampus to long term neocortical stores^5^. Studies thus far have accordingly focused on uncovering evidence for hippocampal involvement in sleep-dependent consolidation for types of memory that depend on the hippocampus for initial encoding^1^.

The hippocampus could conceivably play a broader role, however, by helping to reinstate extra-hippocampal memory traces for processing during sleep. There are hints that this may be the case for a motor learning paradigm known as the motor sequence task (MST)^9^. In this task, participants type a 5-digit sequence (e.g. 4-1-3-2-4) as quickly and accurately as they can for a number of timed trials. Performance on this task has been consistently found to benefit from sleep^10^. The hippocampus seems unlikely to be necessary for performing this task, as there is no long term declarative memory component — the sequence is always displayed for the participant — and the similar serial reaction time task (SRTT) does not require a healthy hippocampus^11,12^ (at least when sequential dependencies are first order^13^).

At the same time, there is evidence that the hippocampus may be involved in offline consolidation of the MST. Hippocampal activity and connectivity with other regions during initial learning is associated with performance improvement across sleep^14-17^, post-learning hippocampal activity during sleep is associated with improvement^18^ and, after sleep, there is increased activity in the hippocampus while performing the task^15,19,20^.

Further indirect evidence that the hippocampus is important for sleep-dependent MST consolidation comes from associations between MST improvement and sleep spindles. Sleep spindles and stage 2 sleep (a stage defined by spindle events) are associated with improvement on the MST^21-32^, and spindles are in turn often associated with hippocampal replay^3-7^. Consistent with the idea that spindles can provide an index of hippocampal involvement in consolidation, they have often been associated with improvement in tasks that are known to depend on the hippocampus^33^ (though not all hippocampally dependent tasks show spindle correlations). In addition, patients with hippocampal sclerosis due to temporal lobe epilepsy and patients with amnestic mild cognitive impairment, which is associated with hippocampal dysfunction^34^, have fewer spindles than normal and deficits in consolidation of hippocampally dependent memory^35,36^.

It is unclear from this collection of findings exactly what role, if any, the hippocampus plays in MST learning and consolidation. First, is the hippocampus required for initial learning? There is extensive neuroimaging evidence that the hippocampus is engaged by performing the MST and other motor learning tasks like the SRTT^17,37-40^, but we cannot determine from neuroimaging studies whether the hippocampus is necessary for normal MST performance or whether it is merely engaged by it. Amnesics’ normal performance on the SRTT^11,12^ favors the latter possibility, though the MST may engage the hippocampus differently. Second, is the hippocampus necessary for normal consolidation of the task? The hippocampus is associated with offline gain in performance in neuroimaging studies, but, again, it is unknown whether this is epiphenomenal.

To answer these questions, we tested learning and consolidation of the MST in four patients with severe amnesia due to hippocampal damage and ten demographically matched control participants. We found that amnesic patients performed similarly to control participants in their initial learning of the MST, indicating that, as in the SRTT, the hippocampus is not required for normal learning. In contrast, unlike controls, the patients exhibited no evidence of sleep-dependent consolidation. These findings are consistent with the prior literature but indicate that the previously observed hippocampal engagement in the initial learning of the MST may be primarily setting the stage for later offline involvement. Our results demonstrate that the hippocampus is necessary for the consolidation of a form of memory that does not require the hippocampus for acquisition, which suggests that the hippocampus plays a broader role in sleep-dependent memory consolidation than was previously understood.

## Results

Four amnesic patients (Figure 1; Table 1) and ten matched control participants trained on 12 trials of the MST on day 1 and were tested on an additional 12 trials 24 hours later on day 2. Each trial consisted of 30 seconds of repeatedly typing a 5-digit sequence “as quickly and accurately as possible” and then resting for 30 seconds. The sequence was continuously displayed on the computer screen in front of the participant as well as on a notecard next to the keypad. Patients completed this two-day protocol twice, using different sequences.

**Figure 1:**
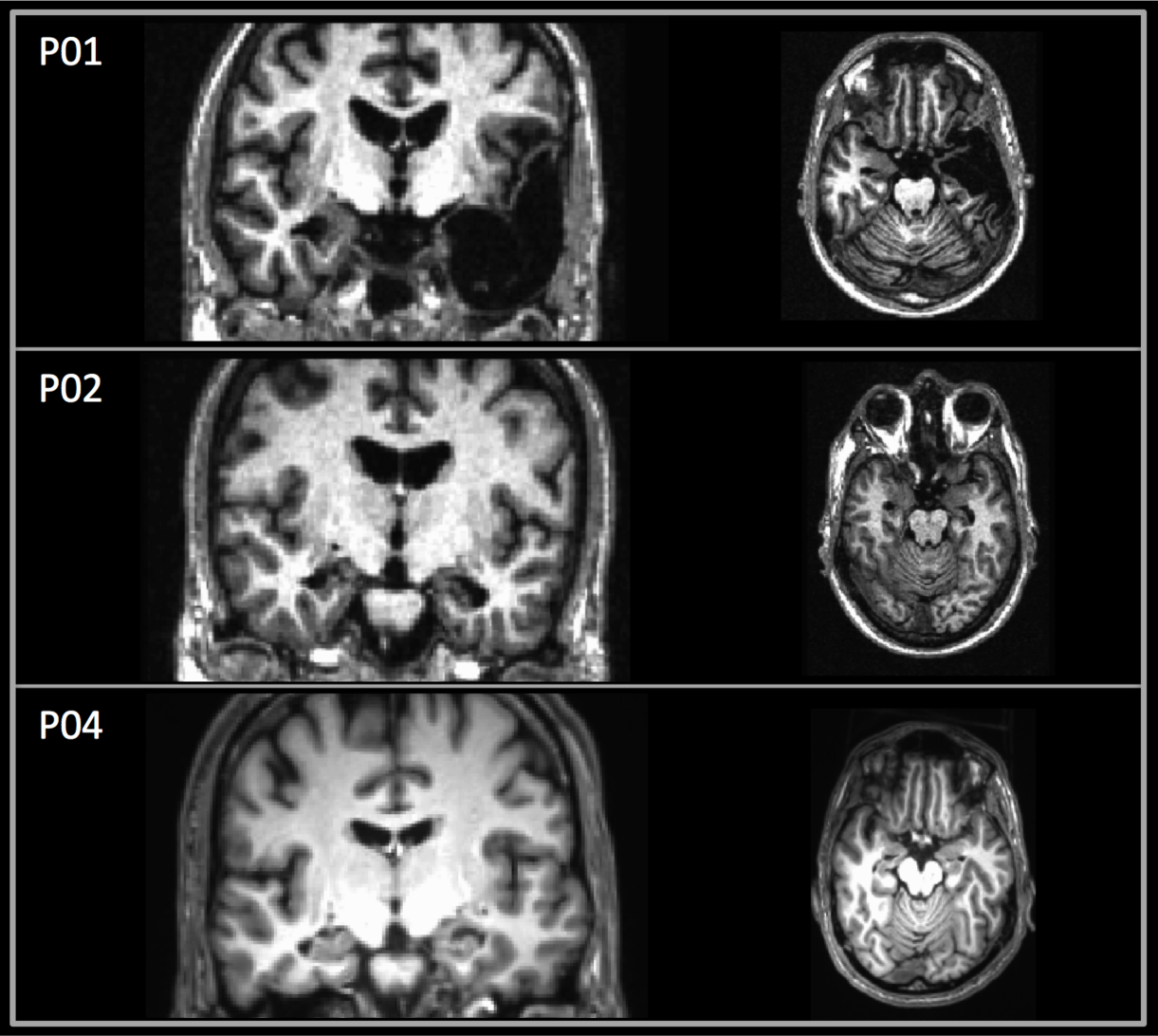
Coronal and axial T1-weighted magnetic resonance images depict lesions for patients P01, P02, and P04 (no scans were available for P03). The left side of the brain is displayed on the right side of the image.

**Table 1.** Demographic and neuropsychological characteristics of amnesic patients.

|  | Etiology | Age | Edu | WAIS III |  | WMS III |  |  | Years since onset | % Volume Loss Hippocampal | % Volume Loss Subhippocampal |
| --- | --- | --- | --- | --- | --- | --- | --- | --- | --- | --- | --- |
|  |  |  |  | VIQ | WMI | GMI | VD | AD |  |  |  |
| P01 | Status epilepticus + left temp. lobectomy | 53 | 16 | 93 | 94 | 49 | 53 | 52 | 27.3 | 63% | 60%<sup>a</sup> |
| P02 | Hypoxic-ischemic | 61 | 14 | 106 | 115 | 59 | 72 | 52 | 24.2 | 22% | --- |
| P03 | Hypoxic-ischemic | 65 | 17 | 131 | 126 | 86 | 78 | 86 | 15.0 | N/A | N/A |
| P04 | Stroke | 53 | 20 | 111 | 99 | 60 | 65 | 58 | 3.45 | 43% | --- |

Note: Age = age in years at time of first training session; Edu = education in years; WAIS-III = Wechsler Adult Intelligence Scale-III; WMS-III = Wechsler Memory Scale-III; VIQ = verbal IQ; WMI = working memory index; GMI = general memory index; VD = visual delayed; AD = auditory delayed. <sup>a</sup>Volume loss in left anterior parahippocampal gyrus (i.e., entorhinal cortex, medial portion of the temporal pole, and the medial portion of perirhinal cortex). See <sup>41</sup> for methodology.

### Sleep and alertness surveys

Controls and patients reported similar amounts of sleep both the night before training (controls = 7.2 ± 1.1 (S.D.) hours; patients = 8.0 ± 1.5; *t*[12]=1.05, *p*=0.32) and the night between training and test (controls = 7.3 ± 1.1 hours, patients = 7.8 ± 1.8; *t*[12]=0.65, *p*=0.53). They also reported similar sleep quality on both nights (1 = slept very poorly to 7 = very well; pre-training: controls = 5.6 ± 1.1, patients = 5.3 ± 0.3; *t*[12]=0.63, *p*=0.54; post-training: controls = 5.3 ± 1.1; patients = 5.8 ± 0.6; *t*[12]=0.78, *p*=0.45) and similar alertness (1 = may fall asleep to 7 = wide awake) both at the time of training (controls = 6.2 ± 0.8, patients = 6.1 ± 0.9; *t*[12]=0.16, *p*=0.88) and at test (controls = 5.9 ± 1.2; patients = 6.3 ± 1.5; *t*[12]=0.46, *p*=0.65).

### Learning time course

Control and patient learning curves were remarkably similar throughout the course of training, whereas the two groups separated in the test phase, with patients performing worse than controls (Figure 2). Both groups displayed a drop in performance from the end of training to the beginning of test and then a quick rise to a stable performance level that was maintained for the rest of test. This pattern has been observed before in older adult participants performing the MST and may reflect a need for older participants to get re-acquainted with the sequence before true performance levels can be expressed^42^. A similar lag in test performance has also been seen in younger subjects performing a 9-digit bimanual version of the task^43^.

**Figure 2:**
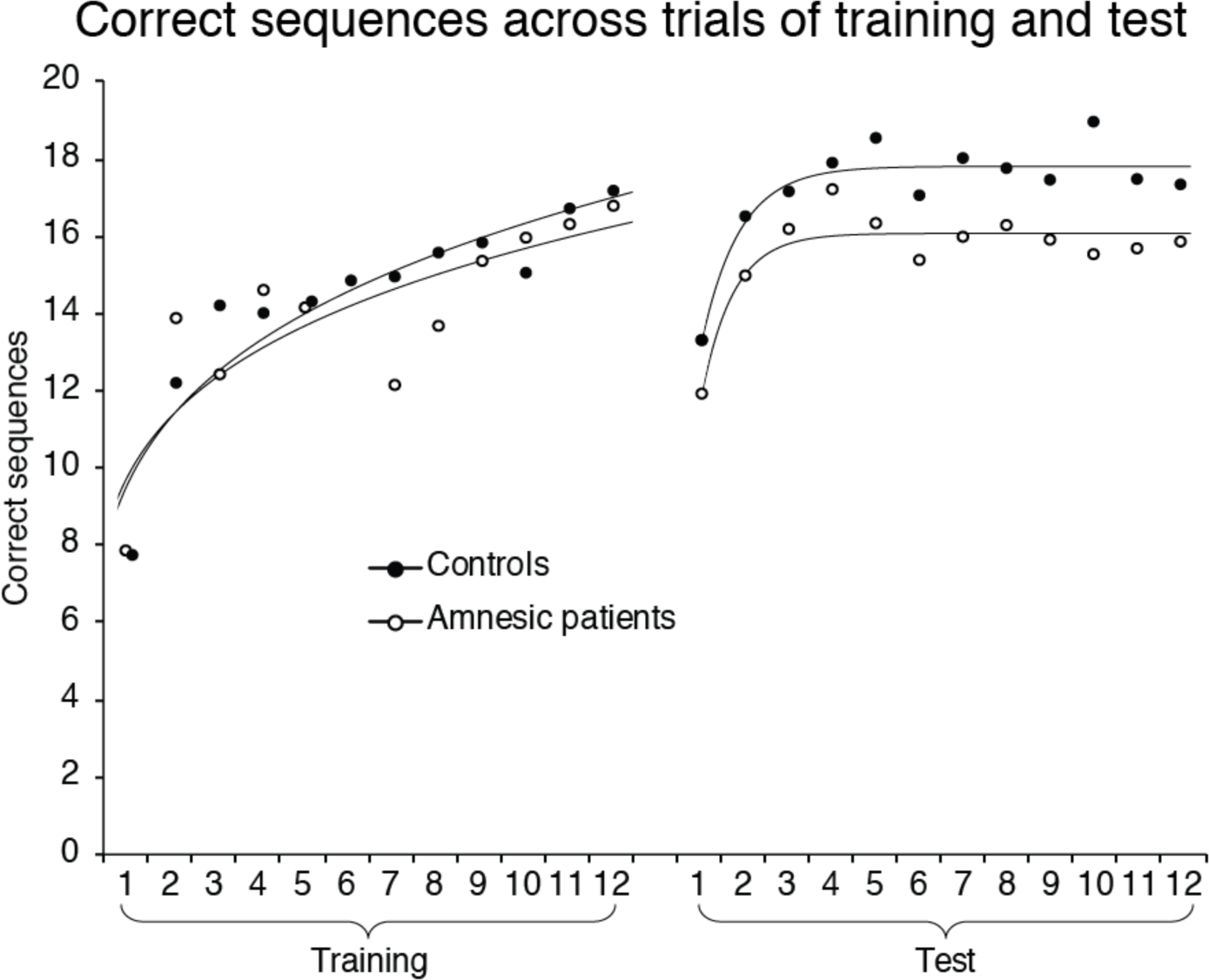
Performance across training and test for controls and amnesic patients. Fitted curves are power functions for training and exponential functions for test.

To test whether group differences were statistically reliable, we ran a mixed effects model. The model revealed reliable effects of day (χ^2^(1)=13.65, *p*=0.0002), trial (χ^2^(1)=78.46, *p*<0.0001), and, critically, a group by day interaction (χ^2^(1)=5.63, *p*=0.018), with the difference between controls and patients larger at test than during training. This indicates that the amnesic patients showed less offline improvement on the MST than controls. There was no effect of sequence (χ^2^(1)=0.06, *p*=0.80), indicating that patients did not perform differently on their first vs. second set of two-day sessions.

### Change from end of training to test

Only the controls showed significant sleep-dependent improvement at test. We calculated for each participant the percent change from the last three training trials to both the first three (initial change) and last six (plateau change) test trials^27^. For the initial change, patients exhibited a near-significant decline in performance (Figure 3; mean= −13.6%, *t*[3]=2.68, *p*=0.075, *Cohen’s d*=1.34) whereas at plateau performance they almost fully recovered to the level of performance achieved at the end of training (mean=−1.9%, *t*[3]=0.36, *p*=0.74, *d*=0.18). Controls, in contrast, performed at the same level at the beginning of test as at the end of training (mean=−0.05%, *t*[9]=0.02, *p*=0.99, *d*=0.01) and showed reliable improvement at plateau (mean=12.4%, *t*[9]=2.54, *p*=0.032, *d*=0.80). The difference between groups was reliable for the initial change (*t*[12]=2.51, *p*=0.027, *d*=1.49; plateau change: *t*[12]=1.68, *p*=0.118, *d*=1.0).

**Figure 3:**
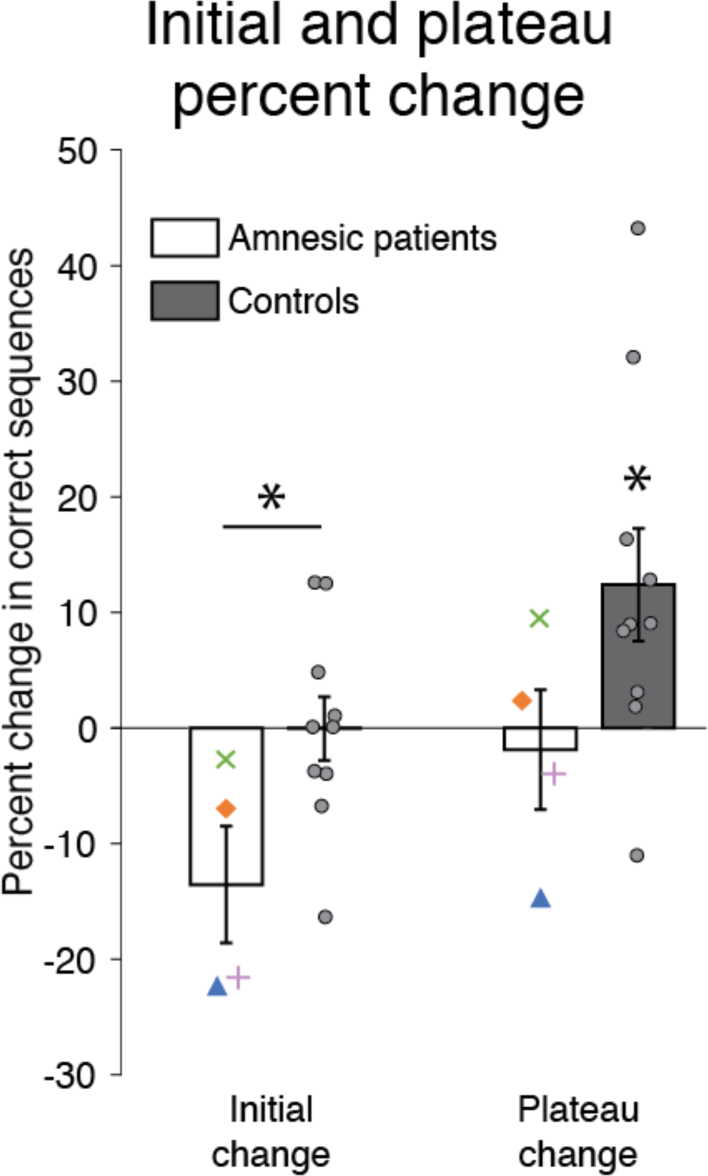
Percent change in performance from training to test. Individual controls are plotted as gray circles. Each patient is plotted with a unique marker: P01 = green ex; P02 = orange diamond; P03 = pink plus; P04 = blue triangle. Asterisk above horizontal line denotes significant difference between groups; asterisk without line indicates where condition differs from zero. Error bars denote ± 1 SEM. * *p*<0.05, t-test.

There was no obvious relationship between degree of hippocampal damage and the consolidation deficits in the patients. P01 had the most extensive damage but showed the least evidence for a consolidation deficit.

### Training performance

To verify that there was no difference in initial learning between the two groups, we assessed the percent change from the first training trial to the last three training trials. Percent improvement across training averaged 131% for patients and 176% for controls, levels which were not significantly different between the groups (*t*[10]=0.41, *p*=0.69, *d*=0.25).

## Discussion

These findings demonstrate a critical role for the hippocampus in the sleep-dependent consolidation of a task that does that require the hippocampus for initial learning. Performance on the MST was assessed in two sessions separated by 24 hours in patients with amnesia due to hippocampal damage and in matched control participants. Patients and controls performed equally well on the first day during training, but patients performed significantly worse than controls on the second relative to first day, and their proportional change in performance from the end of training to the beginning of test was significantly lower than that for controls. The patients had of course retained much of their learning, exhibiting savings in performing the motor sequence across days that has been well documented in amnesics since patient H.M.^44^. But their poor performance at test relative to controls indicates a selective deficit in offline consolidation of the task. Indeed, even at the end of test, patients showed no improvement compared to the end of training. The hippocampus can thus play a critical role in consolidation without being necessary for initial learning.

How might the hippocampus be involved in consolidation of the MST? There is evidence that the hippocampus is engaged during initial learning of the task, and that this activity is related to sleep-dependent consolidation^17^. There are several possibilities for what this activity might reflect. One possibility is that the hippocampus is learning a representation of the sequence in parallel with, and perhaps in interaction with^15^, the striatum. There is extensive evidence that the hippocampus is involved in learning sequential content in both motor^37-40^ and non-motor domains^45,46^ and that it is necessary for normal sequential learning in non-motor domains^47,48^ (perhaps because there is less parallel learning occurring in the striatum in non-motor tasks). The hippocampus may thus benefit consolidation by replaying the content of the sequence during sleep in the same way it would for a paradigm that strictly required the area initially. If the hippocampus is learning the sequential content alongside the striatum, it is likely that this representation would take a different form than the striatal representation. Indeed, there is evidence that the striatum learns the motor contingencies of the MST while the hippocampus learns a more abstract representation of effector-independent spatial contingencies, and that it is this hippocampal version of the representation that undergoes sleep-dependent improvement^10,14,23^.

Another possible role for the hippocampus during initial learning is that instead of learning the content of the sequence per se, it is tagging striatal or motor cortical memories for later offline processing^17,49^. One version of this possibility is that the hippocampus binds incidentally encoded contextual information with sequential representations stored in other areas. This contextual information could then be revisited during sleep, in turn helping to reinstate the sequential representations stored in the other areas. In other words, the hippocampus could be reminding the sleeping brain that it did a sequence learning experiment earlier that day. Future work will be needed to investigate and adjudicate between these possibilities.

We based our protocol on a prior MST study in older adults (ages 60–79) who completed a training session and then a test 24 hours later, including a night of sleep, as in the current study, or a training session in the morning and a test 12 hours later, with no intervening sleep^42^. The participants in the sleep condition showed a similar pattern to our control subjects, with no initial improvement but reliable plateau improvement. The results for participants in the condition with no sleep was remarkably similar to that for our amnesic patients, with worse initial performance and plateau performance at the same level as performance at the end of training. The similarity between the amnesic patients and the healthy older adults with no intervening sleep suggests that the impact of hippocampal damage on sleep-dependent memory processing is similar to that of not having any post-training sleep. This correspondence helps to rule out an alternative explanation for our results, which is that the hippocampus is not needed for consolidation, but instead for reinstating context^50^ of the task from the previous day. If contextual reinstatement by the hippocampus is what accounts for the improved performance in the second session for controls, then participants with a healthy hippocampus who did not sleep between the two sessions in the prior study should have experienced the same benefit. Taking the results from the two studies together, we can conclude that hippocampal damage impairs sleep-dependent consolidation of the MST.

Older adults do not show the same sleep-dependent benefit in initial improvement as younger adults on the MST^21,42,51,52^. Younger adults tend to show more robust consolidation and a boost in performance in the first three trials of the test phase after sleep^28,53^. We followed the same protocol as the study described above that found no improvement in initial test performance in older adults but an improvement in plateau performance^42^, and we replicated those findings here in our control participants. Other studies have looked only at initial performance and found results consistent with ours^19,51^. One additional study found no benefit of a nap for the MST in older adults^21^, which may have been related to poor performance in the training phase^19^. An intriguing possibility is that the difference between younger and older adults on this task may be functionally similar to the difference we observed between patients and controls. There is a known reduction in hippocampal function with healthy aging^54^ as well as a reduction in sleep spindles with age^55^, and the reduction in sleep spindles in older adults has been related to MST consolidation^21^. Thus, young adults, older adults, and patients with amnesia may fall on a spectrum of decreasing contribution of the hippocampus and spindles to consolidation.

There is debate in the literature as to whether the benefit of sleep to the MST for young adults is to boost performance or simply to stabilize it^56,57^, alternatives which tend to support either an active role for sleep or a passive period of rest and reduced interference. Though the present findings do not directly speak to this debate or hinge on it, we believe they are easier to explain from the perspective of an active role for sleep: If a brain region is not critical for initial performance of a task but becomes critical offline, it seems likely that the region is playing an active role during that offline period.

A recent study assessed MST performance in patients experiencing transient global amnesia, a form of hippocampal amnesia lasting less than 24 hours^58^. During training, these patients typed fewer sequences overall than controls but exhibited similar percent improvement across training. Patients were tested on the same sequence again two days later, when they were no longer experiencing amnestic symptoms, and they performed in the same range as control subjects who were learning a new sequence. Improvement across days was larger in patients than controls, which the authors interpreted as evidence that consolidation boosted performance more in patients than controls. However, the findings can also be interpreted simply as better overall performance outside of an acute amnestic state. We therefore do not view these findings as inconsistent with ours. Another recent study assessed learning of the MST in patients with hippocampal dysfunction due to medial temporal lobe epilepsy^59^. During training, patients initially performed at the same level as controls, but their percent improvement across training was reliably lower than controls. This is a counterintuitive finding, as the disruption to hippocampal function in these patients is likely much less than that in our patients, who have substantial hippocampal lesions. One possibility is that hippocampal dysfunction in epilepsy patients may disrupt hippocampal-striatal interactions during learning^17^, whereas larger hippocampal lesions may leave the striatum to act more functionally and independently. This is an interesting possibility to explore in future work. For now, our sample of patients demonstrates that it is possible to exhibit normal MST learning despite extensive hippocampal damage.

While we have demonstrated that the hippocampus can be involved in the consolidation of a task that does not require the hippocampus for initial learning, we are not claiming that the hippocampus is necessarily involved in the offline processing of all tasks that do not initially require the structure. It is conceivable that the hippocampus plays a general enough role in encoding our experience and its context that it will be important for a broad range of tasks that undergo sleep-dependent consolidation, but it is also possible that the hippocampus is involved in consolidation of the MST specifically because it is attuned to sequential information. More work will be needed to determine the precise scope of the region’s involvement. Our findings open the door to these new possibilities by providing a proof of concept that the hippocampus can be critical offline for a task for which it was not necessary during initial learning.

## Methods

### Participants

Eight patients with medial temporal lobe lesions (6 males, 7 right-handed) and 12 control participants (10 males, 8 right-handed) participated in the study. Etiology for the patients was hypoxic-ischemic injury secondary to cardiac or respiratory arrest (n=5), encephalitis (n=1), stroke (n=1), and status epileptics followed by left temporal lobectomy (n=1). All patients were in the chronic phase of illness, with time post injury ranging from 3.5 to 36.4 years (mean=21.1).

Four patients and two controls did not meet the inclusion criterion on their first session, which required a minimum of 10 correct sequences on average over the last three trials of training. The average scores for the excluded patients were 8.6, 3.7, 3.2, and 3.1 sequences. The patient with the highest score was tested on a second sequence, but again failed to meet threshold, with a score of 9.3. The other three patients were not tested on a second sequence. We do not believe that these low scores reflect a sequence learning deficit, but rather a motor deficit. These four patients were also the slowest of all participants on the warmup task (described below), which does not require sequence learning. Two of these subjects had basal ganglia damage: one had extensive volume reduction in caudate, putamen, and pallidum bilaterally, and the other had reduction in left pallidum only. It is possible that this damage contributed to the slower motor performance in these patients.

Demographic and neuropsychological characteristics for the four included patients are provided in Table 1. The neuropsychological profiles of each patient indicated severe episodic memory impairment (mean General Memory Index = 64.5), with otherwise preserved cognition (mean VIQ = 110.3; mean Working Memory index = 108.5). Lesions for three of the patients are shown in Figure 1. The remaining patient (P03) had suffered cardiac arrest, and could not be scanned due to medical contraindications. Medial temporal lobe pathology for this patient was inferred based on etiology and neuropsychological profile. Two patients (P02 and P04) had lesions restricted to the hippocampus, and one patient had volume loss extending outside of the hippocampus (P01).

The 10 control participants included in analyses were well matched to the included patients in terms of sex (8 males; all included patients male), handedness (9 right-handed; all included patients right-handed), age (mean=57.7; patients mean=58.0), years of education (mean=14.7; patients mean=16.8), and VIQ (mean=112.1; patients mean=110.3).

All participants provided informed consent in accordance with the Institutional Review Board of VA Boston Healthcare System.

### Procedure

The task was presented to participants on a laptop using MATLAB with Psychophysics Toolbox^60^. Participants were instructed to rest four fingers of their left hand on a button box with buttons labeled 1, 2, 3, and 4. At the beginning of each session, participants completed a warmup task, where they were instructed to repeatedly type the sequence 1-2-3-4 when the screen turned from red to green and to “try to be as fast and accurate as you can.” The sequence was always displayed on the screen, during both rest and typing periods. The number of seconds until the screen turned green was then displayed as spelled out numbers (“ten”, “nine”, “eight”…). The screen remained green for 30 seconds. With every key press, a new dot appeared in a horizontal line on the screen, to provide feedback to the participant that their key press was registered. After the line reached the right side of the screen, the dots then disappeared one at a time with each additional key press. The experimenter wore one ear bud through which she would hear a beep whenever the participant pressed a button out of sequence. This allowed her to provide rapid feedback if the participant was not pressing the buttons correctly.

After 30 seconds, the screen turned red, and participants were instructed to stop typing and take a 30 second break. A thirty second countdown immediately began with the number of seconds left again spelled out on the screen. At the end of the countdown, the screen turned green again for 30 seconds, and the participants again typed the warmup sequence. If the participant did not yet seem comfortable with the task, the experimenter had the option to initiate additional warmup trials.

Once the participant was accustomed to the task, the experimenter initiated the training phase. The training had the same structure as the warmup, with 30 seconds of typing interspersed with 30 seconds of rest. One of four sequences was used: 4-1-3-2-4, 1-4-2-3-1, 3-1-4-2-3, or 2-4-1-3-2. The sequence assignment was counterbalanced across subjects. In addition to the sequence being displayed continuously on the screen, an index card displaying the sequence was also placed next to the keypad so that participants did not have to look at the screen while typing. Participants completed 12 trials of training. Throughout the session, the experimenter reminded participants to start and stop typing as needed.

24 hours later, participants completed another warmup and then the test, which consisted of 12 identical trials of the same sequence that they had typed the previous day. In order to obtain better estimates of each patient’s performance, patients who met the performance criterion (10 sequences correct across the last three trials of training) for their first sequence were tested again several months later on a second sequence. The sequence assignment was again counterbalanced across participants. Patients completed the same two-day procedure with the second sequence.

On each day of training or testing, participants filled out a survey asking how well they slept the previous night, the duration of their sleep, and how alert they felt. On the morning of test days, participants also answered these questions using a paper survey filled out at home around the time of awakening.

### Mixed effects model

To assess whether the patients and controls differed in their behavior across the training and test days, we fit a mixed effects model, with participant as random effect and group, day, trial, and sequence as fixed effects:

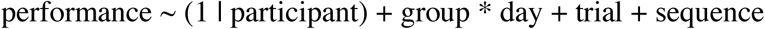
where day indicates training vs. test, and sequence indicates the first or second sequence used for patients. Significance of factors was assessed by removing the factor (or interaction between factors) from the model and assessing the difference between the original and modified models using the χ^2^ statistic.

### Trial outlier removal

As a preprocessing step for percent change analyses, where a small subset of trials was used (making the analyses potentially sensitive to trial outliers), we removed trials for each participant that fell far from an estimated learning curve, as follows. For each participant and each day, a power function (y=b•x^m^) was fit to performance across the 12 trials. The squared residuals for each trial were calculated, and trials falling more than 2 SD outside the distribution of squared residuals across all trials and subjects were excluded. This resulted in exclusion of 19 trials out of 432 (4.4%). 12 of these trials came from patients and 7 from controls. We then averaged the data across the two sequences that each patient completed. When a trial was missing from one sequence and not the other, the non-missing data point was used. This resulted in just one missing trial across all the patient data. Supplementary Figure 1 shows the data for individual controls and patients after this trial exclusion process (dashed blue line indicates the one missing trial for patient data). These preprocessing steps served to provide smoother estimates of patient performance given the relatively small number of patients, minimizing the influence of outlier trials.

### Percent change calculation

Our estimate of the “initial” percent change from the end of training to the beginning of test was 100•(mean of first 3 test trials–mean of last 3 training trials)/mean of last 3 training trials. The “plateau” change was calculated as 100•(mean of last 6 test trials–mean of last 3 training trials)/mean of last 3 training trials. Percent improvement over the course of training was 100•(mean of last 3 training trials– first training trial)/first training trial. Two control subjects were not included in this analysis: for one, the first trial of training was excluded as an outlier, and for the second, a technical issue resulted in loss of data for the first three trials of training (this was the only data loss that occurred during the study). Differences between groups and differences of each group from zero were calculated using two-tailed t tests.

### Data Availability

Behavioral data will be made available with manuscript publication.

## Acknowledgments

We thank James Antony for helpful discussions. This work was supported by: NIH F32-NS093901 (ACS); NIH R01-MH48832 (RS); NIH R01-MH67720 (DSM); NIH K24-MH099421 (DSM); Senior Research Career Scientist Award from the Clinical Science Research and Development Service, Department of Veterans Affairs (MV). The contents of this manuscript do not represent the view of the US Department of Veterans Affairs or the US Government.

